# Neurocan regulates axon initial segment organization and neuronal activity in cultured cortical neurons

**DOI:** 10.1101/2024.01.26.577401

**Authors:** David Baidoe-Ansah, Hadi Mirzapourdelavar, Stepan Aleshin, Björn H Schott, Constanze Seidenbecher, Rahul Kaushik, Alexander Dityatev

## Abstract

The neural extracellular matrix (ECM) accumulates in the form of perineuronal nets (PNNs), particularly around fast-spiking GABAergic interneurons in the cortex and hippocampus, but also in association with the axon initial segments (AIS) and nodes of Ranvier. Increasing evidence highlights the role of Neurocan (Ncan), a brain-specific component of ECM, in the pathophysiology of neuropsychiatric disorders like bipolar disorder and schizophrenia. Ncan localizes at PNNs, nodes of Ranvier and the AIS, highlighting its potential role in regulation of axonal excitability. Here, we used knockdown and knockout approaches in mouse primary cortical neurons in combination with immunocytochemistry, western blotting and electrophysiological techniques to characterize the role of Ncan in the organization of PNNs and AIS and in the regulation of neuronal activity. We found that reduced Ncan levels led to remodeling of PNNs around neurons via upregulated Aggrecan mRNA and protein levels, increased expression of activity-dependent c-Fos and FosB genes and elevated spontaneous synaptic activity. The latter correlated with increased levels of Ankyrin-G in the AIS particularly in excitatory neurons, and with the elevated expression of Na_v_1.6 channels. Our results suggest that Ncan regulate expression of key proteins in PNNs and AISs and provide new insights into its role in the fine-tuning of neuronal functions.

## INTRODUCTION

Neuronal excitability and synaptic communication in the central nervous system (CNS) rely on the initiation of action potentials (APs) at the axon initial segment (AIS) (Huang and Rasband, 2018; Torii et al., 2020) and its propagation along the axon towards the synaptic terminals. The initiation and propagation of APs are facilitated by the aggregation of voltage-gated sodium (Na_v_) and potassium channels (Leterrier et al., 2017). Interestingly, Na_v_ channels cluster at densities shown to be more than 50-fold higher in the AIS as compared to dendrites (Lorincz & Nusser, 2010a). At the AIS, different Na_v_ isoforms have been identified, including Na_v_1.1, Na_v_1.2, and Na_v_1.6 (Battefeld et al., 2014; Lopez et al., 2017), which serve as the key ion channels in the regulation of neuronal excitability. These channels are distributed at distinct locations along the AIS, with the proximal region mostly containing both Na_v_1.1 and Na_v_1.2 whereas Na_v_1.6 are enriched at the distal AIS and at the node of Ranvier (Freeman et al., 2016; Wang et al., 2017). The clustering of Na_v_1.2 and Na_v_1.6 is regulated differently (Kaplan et al., 1997), and they also differ functionally. While Na_v_1.6 initiates APs in both excitatory and inhibitory neurons (Wang et al., 2017) Na_v_1.2 controls AP transmission, especially the backpropagation of APs (Wang et al., 2017). The clustering and distribution of Na_v_ depend on the adaptor protein Ankyrin G (AnkG) (Brachet et al., 2010). Structurally, AnkG regulates the localization and diffusion (Bender & Trussell, 2012; Lorincz & Nusser, 2010b) of ion channels as well as other AIS proteins including the cell adhesion molecule Neurofascin-186 and βIV-spectrin (Hedstrom et al., 2007; Leterrier et al., 2015).

The AIS is also enriched in some extracellular matrix (ECM) molecules (Bekku & Oohashi, 2010; Bruckner et al., 2006; Dityatev, Seidenbecher, et al., 2010). The neural ECM generally comprises hyaluronic acid, proteoglycans, and a variety of other glycoproteins (Dityatev & Rusakov, 2011), and it exists in diffuse forms as perisynaptic and interstitial ECM or in more dense forms as periaxonal coats and perineuronal nets (PNN). PNNs mostly surround inhibitory neuronal cell bodies and proximal dendrites (Dityatev & Schachner, 2006) and are made up of chondroitin sulfate proteoglycans (CSPGs) attached to the hyaluronic acid backbone, stabilized by link proteins and the glycoprotein Tenascin-R (TnR) (Lau et al., 2013). CSPGs found in PNNs are aggrecan (Acan), brevican (Bcan), neurocan (Ncan), and versican (Vcan) which also occur in varying degrees of expression at the AIS (Hedstrom et al., 2007). Bcan, Vcan, and TnR are highly enriched at the AIS whereas Acan and Ncan are expressed in a lesser amount (Frischknecht et al., 2014; Hedstrom et al., 2007; Susuki et al., 2013). The AIS ECM is more solid than PNNs having a mesh-like appearance with “holes”, and it can be more broadly found in association with both PNN-positive and -negative neurons (Hedstrom et al., 2007; John et al., 2006).

Several studies have explored the role of neural ECM composition in neuronal excitability. For instance, digestion of chondroitin sulfates with chondroitinase ABC (chABC) modulates the excitability of parvalbumin-expressing interneurons (Dityatev et al., 2007; Hayani et al., 2018). In contrast, the digestion of heparan sulphates with heparinase reduces the excitability of excitatory neurons and downregulates AnkG at the AIS (Minge et al., 2017) in a calcium/calmodulin kinase II (CaMKII)-dependent manner (Song et al., 2023). Knockdown of Bcan in PV+ interneurons reduced their excitability via modifying clustering of potassium channels (Favuzzi et al., 2017). These studies suggest that ECM molecules are essential for regulation of neuronal excitability. Ncan has recently received considerable attention as the human NCAN gene has been identified as a genetic risk factor for bipolar disorder and particularly the manic phenotype (Cichon et al., 2011; Miro et al., 2012), and for schizophrenia (Mühleisen et al., 2012). Furthermore, data from animal models and patients show dysregulation of Ncan expression in several CNS disorders associated with abnormalities in neuronal and network activity, such as epilepsy (Okamoto et al., 2003), depression (Yu et al., 2022), or Alzheimer’s disease (Yan et al., 2016). In healthy humans, the NCAN psychiatric risk variant rs1064395 has been associated with hippocampus-dependent memory function and prefrontal gray matter density (Assmann et al., 2021; Dannlowski et al., 2015; Raum et al., 2015), but the mechanistic role played by Ncan has not been studied.

Thus, in this study, we used both knockout and knockdown approaches to elucidate the roles of Ncan in the organization of PNNs and AISs and in the regulation of neuronal activity. The knockdown approach was employed to avoid activation of the potential genetic compensatory mechanisms common for knockout models (El-Brolosy & Stainier, 2017; Zimmer et al., 2019). As experimental system, we used dissociated mouse primary cortical neurons from Ncan KO and wildtype mice. We knocked down Ncan using AAVs to express one of two Ncan shRNAs in mouse primary cortical neurons from DIV 7 to DIV 21 and investigated the effects of Ncan depletion on the expression of various ECM components of PNNs and AISs using RT-qPCR, Western blotting and immunocytochemistry. Additionally, we used ICC and whole-cell patch-clamp recordings to unveil the effect of Ncan depletion on AnkG levels and Nav levels and neuronal activity.

## METHODS

### Knockdown experiments using Ncan shRNA

To knock down mouse Ncan (GeneID: NM_007789.3), shRNA plasmids were cloned by the insertion of the siRNA sequences (Sigma) shown to be effective against Ncan mRNA (Okuda et al., 2014; Su et al., 2017) (Suppl. Table 1) targeting the open reading frame into AAV U6 GFP (Cell Biolabs Inc., San Diego, CA, USA) using BamH1 (New England Biolabs, Frankfurt a.M., Germany) and EcoR1 (New England Biolabs, Frankfurt a.M., Germany) restriction sites. Two shRNAs were produced targeting Ncan messenger RNA (mRNA) (Ncan shRNA) by selecting target sequences in the coding regions of Ncan to prevent or minimize off-target effects (Taxman et al., 2006). The shRNA universal negative control (Sigma, Control shRNA) was used as a non-targeting control (Suppl. Table 1). Positive clones were sequenced and used for the production of recombinant adeno-associated particles as described previously (Mitlöhner et al., 2020). HEK 293T cells were transfected using PEI (1ng/ul) with an equimolar mixture of the shRNA encoding AAV U6 GFP, pHelper (Cell Biolabs Inc., San Diego, CA, USA), and RapCap DJ plasmids (Cell Biolabs Inc., San Diego, CA, USA). From 48-72 hours after transfection, freeze-thaw cycles were implemented to lyse cells and then with benzonase (50 U/mL; Merck Millipore, Burlington, MA, USA) for 1 h at 37 °C. Lysates were centrifuged at 8000× g at 4 °C, supernatants collected, and filtered with a 0.2-micron filter. We then purified filtrates using the pre-equilibrated HiTrap Heparin HP affinity columns (GE HealthCare, Chicago, IL, USA), followed by washing with the following buffers in sequence; Wash Buffer 1 (20 mM Tris, 100 mM NaCl, pH 8.0; sterile filtered) and wash buffer 52 (20 Mm Tris, 250 mM NaCl, pH 8.0; sterile filtered). Elution buffer (20 mM Tris, 500 mM NaCl, pH 8.0; sterile filtered) was then used to elute viral particles. Finally, Amicon Ultra-4 centrifugal filters with 100 kDa molecular weight cutoff (Merck Millipore, Burlington, MA, USA) along with 0.22 μM Nalgene® syringe filter units (sterile, PSE, Sigma-Aldrich, St. Louis, MO, USA) were used to further purify viral particles before being aliquoted and stored at −80 °C.

### Cell culture, neuronal infection, RT-qPCR, and GI-LTP

Cortical neurons were isolated from embryonic (E18) C57BL6/J mice or Ncan KO mice (Zhou et al., 2001), as described previously (Seibenhener & Wooten, 2012). After harvesting, neurons were plated on poly-l-lysine-coated (Sigma; Cat. No. 2636) in 6 well plates without coverslips at a cell density of 200,000 (hippocampal neurons) and 500,000 (cortical neurons) per well. Neurons were maintained in 2.5ml of Neurobasal media (Invitrogen) supplemented with 2% B27 and 1% L-glutamine and 1% Pen-Strep (Life Technologies). Dissociated wildtype and Ncan KO cortical neurons were initially infected with Control and the two Ncan shRNA AAVs at 7 days in vitro (DIV) and mRNA expression levels of Ncan were assayed at DIV-21. Knockdown efficiencies of the two Ncan shRNAs normalized to the Control shRNA were determined using RT-qPCR. From cortical cultures, total RNA was extracted using the EURx GeneMatrix DNA/RNA Extracol kit (Roboklon, Cat. No. E3750) according to the manufacturer’s recommendations (Baidoe-Ansah et al., 2022) and products were further checked for genomic DNA contamination by using Nano-drop and gel electrophoresis to measure the RNA yield, purity, and integrity, respectively. Then, 2 μg of RNA was used for cDNA conversion using the High-Capacity cDNA Reverse Transcription Kit (Cat. No. 4368814). qPCR analysis was performed with TaqMan™ gene expression array assay (ThermoFisher Scientific, Cat. No. 4331182) using the Quant-Studio-5 (Applied Biosystems). Details of all TaqMan™ probes used are given in (Suppl. Table 2). Finally, gene expression per sample was normalized relative to the expression of glyceraldehyde 3-phosphate dehydrogenase (GAPDH) (Meldgaard et al., 2006). For the glycine-induced form of chemical LTP (GI-LTP), already shRNA1 AAV infected cortical cultures were first incubated with sterile artificial cerebrospinal fluid (ACSF) containing the following (in mM): 119.0 NaCl, 1.3 MgSO_4_ 1.0 NaH_2_PO_4_, 26.2 NaHCO_3_, 2.5 KCl, 2.5 CaCl_2_, 11.0 glucose, 0.2 glycine, for 5 minutes at 37°C before returning to ACSF without glycine for 50 minutes (Fortin et al., 2010), whereas control cultures were incubated in ACSF.

### Immunocytochemistry (ICC)

Cortical neurons from wildtype and Ncan KO mice from DIV 21-23 were used for immunocytochemistry experiments. Neurons were washed and fixed with phosphate-buffered saline (PBS) and 4% paraformaldehyde (PFA) for 10 minutes, respectively. They were then permeabilized with 0.1% Triton-X-100 in PBS for 10 min, washed 3 times, and blocked (0.1% Glycine + 0.1% Tween-20 + 10% Normal Goat Serum in PBS) for 60 min at room temperature. Then, the neurons were labeled with primary and secondary antibodies (Suppl. Table 3 and 4, respectively), and finally mounted (Fluoromount; Sigma Aldrich, F4680-25ML) for imaging.

### Imaging and Data acquisition

Next, the mounted coverslips were imaged using a Zeiss LSM 700 confocal microscope. Z-stacked images of neurons were randomly obtained across the entire surface of the coverslips, barring the edges, using a 63x/1.4 NA oil immersion objective. Throughout all the imaging sessions, we maintained the acquiring conditions to compare fluorescence intensity between groups. For analysis of AnkG images, we used the previously described procedure (Minge et al., 2017). Here, using GFP signal to identify shRNA infected neurons and the AnkG immunolabeling, the AISs were first manually outlined and analyzed from the soma edge over a 40 µm long distance with a line profile (width=3.0) with Fiji (Minge et al., 2017). For the quantification of ECM molecules in shRNA-treated cortical neurons, we automatically outlined the profile of the cell somata of GFP+ and Acan+ neurons and created a band of 1.5 μm as the region of interest (ROI). Additionally, for FosB images, we used GFP+neurons to automatically select soma of neurons, then we masked and overlayed the masks over FosB original images to measure fluorescent intensity as well as percentage density.

### Further processing of AnkG data

The experimentally obtained distribution of AnkG along the length of the axon was binned with a one-micron step size. The resulting data were averaged for each group under investigation. The resulting curve was fitted with the sum of two exponentials with weights A1 and A2 plus a baseline (b):

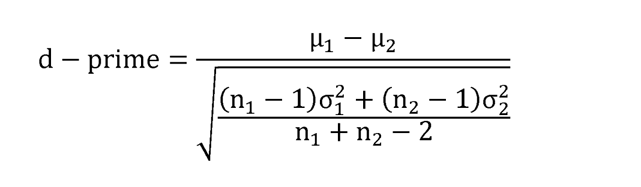

Subsequently, numerical values were obtained for each fitted curve: the amount of AnkG at the beginning of the axon (Y0), the peak height (Y peak) and position (X peak), as well as the distance from the peak where the AnkG amount decreased by a factor of two (T decay).

To estimate the confidence intervals of the parameters, bootstrap resampling was employed. Specifically, for each group under investigation, a sample of axons corresponding to that group was randomly selected with replacement. The described parameters were then computed based on the resampled data. This procedure was repeated 10_4_ times, and the standard deviations of the resulting distributions were represented on the figures as error bars.

Statistical significance of pairwise comparisons was assessed using shuffling. In detail, when comparing two groups, a subset of experimental data containing axons exclusively from one of the groups was extracted. The axon’s group membership was then randomly assigned, and the data were averaged to compute the parameters of the AnkG distribution along the axon for each of the two pseudogroups. After 10_4_ repetitions, the null distribution for each parameter was obtained as the difference between the distributions for each group. The p-value was calculated as the quantile corresponding to the observed difference between the parameters.

### SDS-PAGE and Western blotting

The GeneMATRIX DNA/RNA/Protein Purification Kit (EURX Cat. No. E3597) was used to purify total protein from cortical cells. The concentration of the resulting protein pellet was determined using the Pierce™ BCA Protein Assay Kit (ThermoFisher Cat. 23225) and a NanoDrop™ 2000/2000c spectrophotometer (ThermoFisher). The protein samples were boiled for 5 minutes at 95°C and 10 µg were loaded onto each lane of a 10% SDS gel. Electrophoresis was carried out for 2 hours at 90 V in a tank containing cold 1X TGS (Tris/Glycine/SDS) buffer. The separated proteins were transferred onto nitrocellulose membranes using the Trans-Blot Turbo transfer system (Bio-Rad). To block nonspecific binding, a tris-buffered saline solution with 0.1% tween20 (TBST) supplemented with 5% skim milk (SERVA) was used for 1 hour at room temperature. The primary antibodies were diluted in TBST/1% milk and applied to the membranes overnight on a shaker at 4°C. The blots were washed three times for 10 minutes with TBST and incubated for one hour with Horseradish Peroxidase (HRP)-conjugated secondary antibodies diluted at 1/10,000. After three washes in TBST, the blots were developed using the SuperSignal® West Pico Chemiluminescent Substrate (ThermoScientific Cat. 34078). The membranes were incubated with a working solution containing peroxide solution and luminol enhancer at a 1:1 ratio for 5 minutes at room temperature. The excess reagent was removed and the blots were imaged using a chemiluminescence Imager (Azure Biosystems C300) and covered with clear plastic wrap.

### Whole-cell patch-clamp recording

Recording spontaneous excitatory postsynaptic currents (sEPSCs) allows for the estimation of overall neuronal activity in the absence of stimulation when the membrane potential is held at resting potential (-70 mv) using voltage-clamp technique. Whole-cell patch clamp recordings were performed as described previously (Handara et al., 2019). Briefly, coverslips containing cortical cells were immediately transferred from culture media to the submerged recording chamber and immersed in pre-warmed recording solution containing (in mM): 105 NaCl, 3 KCl, 10 HEPES, 5 Glucose, 2 CaCl_2_, and 1 MgCl_2_. Recordings were obtained under visual control using infrared differential interference contrast optics (Slicescope, Scientifica, UK) and 40X water immersion objective on the SliceScope Pro 6000 upright fixed-stage microscope (Scientifica) and a patch-clamp EPC10 amplifier (HEKA, Lambrecht, Germany). The patch pipettes were pulled from borosilicate glass capillary (wall thickness 0.315 mm, length 100 mm, outer diameter 1.5 mm, Hilgenberg) using a DMZ universal electrode puller (Zeitz Instruments GmbH, Martinsried, Germany) and filled with intracellular solution containing (in mM): 90 KCl, 3 NaCl, 5 EGTA, 5 HEPES, 5 Glucose, 0.5 CaCl_2_, and 4 MgCl_2_. A 3-5 MΩ resistance has been achieved when the pipets are filled with intracellular solution. The whole-cell configuration was obtained by providing a brief suction after establishing a MΩ seal. Only recordings were approved if the series resistance was less than 20 MΩ and did not fluctuate by more than 20% during the experiment. The holding potential was adjusted to -70 mV and the recordings were filtered at 10 kHz and sampled at 20 kHz. Properties of sEPSC were calculated by MiniAnalysis (Synaptosoft Inc., USA) using detection parameters of 5 pA for event threshold. To analyze the characteristics of neuronal activity, we developed a method to detect bursts in sEPSC recordings. First, the moving median absolute deviation of the sEPSC signal using a 100 ms window was calculated and the resulting curve was binarized using a threshold of 20 pA and applied a median filter with a 100 ms window to smooth the signal. Any periods below the binarized signal were considered as no-burst intervals. We then excluded non-burst intervals shorter than 0.2 seconds and identified the start and stop times of the remaining non-burst intervals. To further refine our burst analysis, we excluded any bursts that were shorter than 0.1 seconds and had a maximal or average amplitude below 250 pA and 50 pA, respectively. To avoid contamination of non-burst intervals by bursts, we defined a “protection” zone around each burst as a time window that extended 50 ms before and 200 ms after the burst. Events within this time window were considered “protected” events and were excluded from the analysis.

The final output of our method was a classification of each time point in the original sEPSC signal. Time points within bursts were classified as “bursty”, time points within the protection zone were classified as “protected”, and time points outside of bursts and the protection zone were classified as “nonbursty”. Protected events were excluded from the analysis to ensure that non-burst intervals were not contaminated by bursts.

### Statistical analysis

Statistical analysis was performed using GraphPad Prism 8.0 (GraphPad Software Inc., La Jolla, USA) and Statistica 8.0 (StatSoft, USA) software. All data are shown as mean ± SEM with n being the number of mice. Asterisks in figures indicate statistical significance (with details in the figure legend or Results). The hypothesis that experimental distributions follow the Gaussian law was verified using Kolmogorov-Smirnov, Shapiro-Wilk, or D’Agostinio tests. For pairwise comparisons, we performed the Students’ t-test where the samples qualify for the normality test; otherwise, the Mann-Whitney test was employed. Additionally, Wilcoxon matched-pairs test was used for paired data that did not pass the normality test. Holm-Sidak’s multiple comparisons t-test was used for independent comparisons. Spearman correlation coefficients were computed to estimate the correlation between the variables studied. As indicated, one and two-way ANOVA with uncorrected Fisher’s LSD as well as Brown-Forsythe with Welch ANOVA tests, were also used. The p-values represent the level of significance as indicated in figures by asterisks (*p < 0.05, **p < 0.01, ***p < 0.001 and **** p < 0.0001).

## RESULTS

In this study, we used both knockdown and knockout approaches to elucidate the role of Ncan in neuronal excitability control. As experimental system, we employed dissociated mouse primary cortical neurons from Ncan KO and wildtype mice, which were infected with either Control shRNA or one of two Ncan shRNAs.

### Ncan depletion results in dysreguled expression of PNN components in vitro

To thoroughly dissect Ncan functions, we studied the effects of attenuated Ncan expression *in vitro* by generating 2 AAVs encoding shRNAs targeting the Ncan mRNA and evaluated fold changes in mRNA and protein levels by RT-qPCR and ICC, respectively (Fig. 1 A). Quantification of Ncan expression was conducted in dissociated cortical neurons infected with Ncan shRNA, control vectors (Ctrl), and Ncan KO neurons that were infected with control vectors (Ncan KO) at DIV7 for a duration of 14 days. Following normalization to the internal control, *Gapdh*, our findings indicated that shRNA1 led to an approximate 50% reduction in *Ncan* expression, while shRNA2 resulted in a more prominent 90% reduction compared to the Ctrl condition (Fig. 1B). As expected, *Ncan* transcripts were not detectable in Ncan KO neurons (Fig. 1B). Subsequently, we examined the specificity of our constructs and their impact on the expression of other PNN molecules. Notably, the reduction of *Ncan* was associated with upregulated *Acan* mRNA levels in both shRNA1-and shRNA2 expressing groups, as well as in Ncan KO neurons (Fig. 1B). However, no significant difference was observed in the mRNA levels of another PNN proteoglycan, Bcan. Moreover, a reduced expression of *Hapln1* (Hyaluronan and Proteoglycan Link Protein 1), was observed in shRNA1 expressing neurons, while *Hapln1* expression was upregulated in both shRNA2 and Ncan KO neurons (Fig. 1B). Intriguingly, reduced *Ncan* levels were found to be associated with a corresponding decrease in mRNA levels of *Hapln4*. Conversely, Ncan KO neurons exhibited an upregulation of *Hapln4* mRNA levels (Fig. 1B). To confirm these findings, complementary protein quantification for Acan was done in shRNA1-and shRNA2-treated as well as in Ncan KO neurons compared to Ctrl (Fig. 1C,D). Interestingly, Acan levels increased significantly in the shRNA1 and shRNA2 groups relative to Ctrl in both western blot and ICC analyses (Fig. 1 D, E, F). Expression analysis for two developmentally regulated genes, *Pv* and *Kcnc1*, which are selectively expressed in high-frequency interneurons associated with PNNs, revealed a decreased expression in shRNA1-and shRNA2-expressing neurons, while the expression levels in Ncan KO neurons were similar to Ctrl (Fig. 2A). Analysis for the neuronal activity markers *c-Fos* and *FosB* showed elevated levels in all treated groups compared to the Ctrl group (Fig. 2B). Consistent with mRNA levels, the protein level of FosB was increased, suggesting elevated overall neuronal activity following Ncan depletion (Fig. 2C).

**Figure 1.**
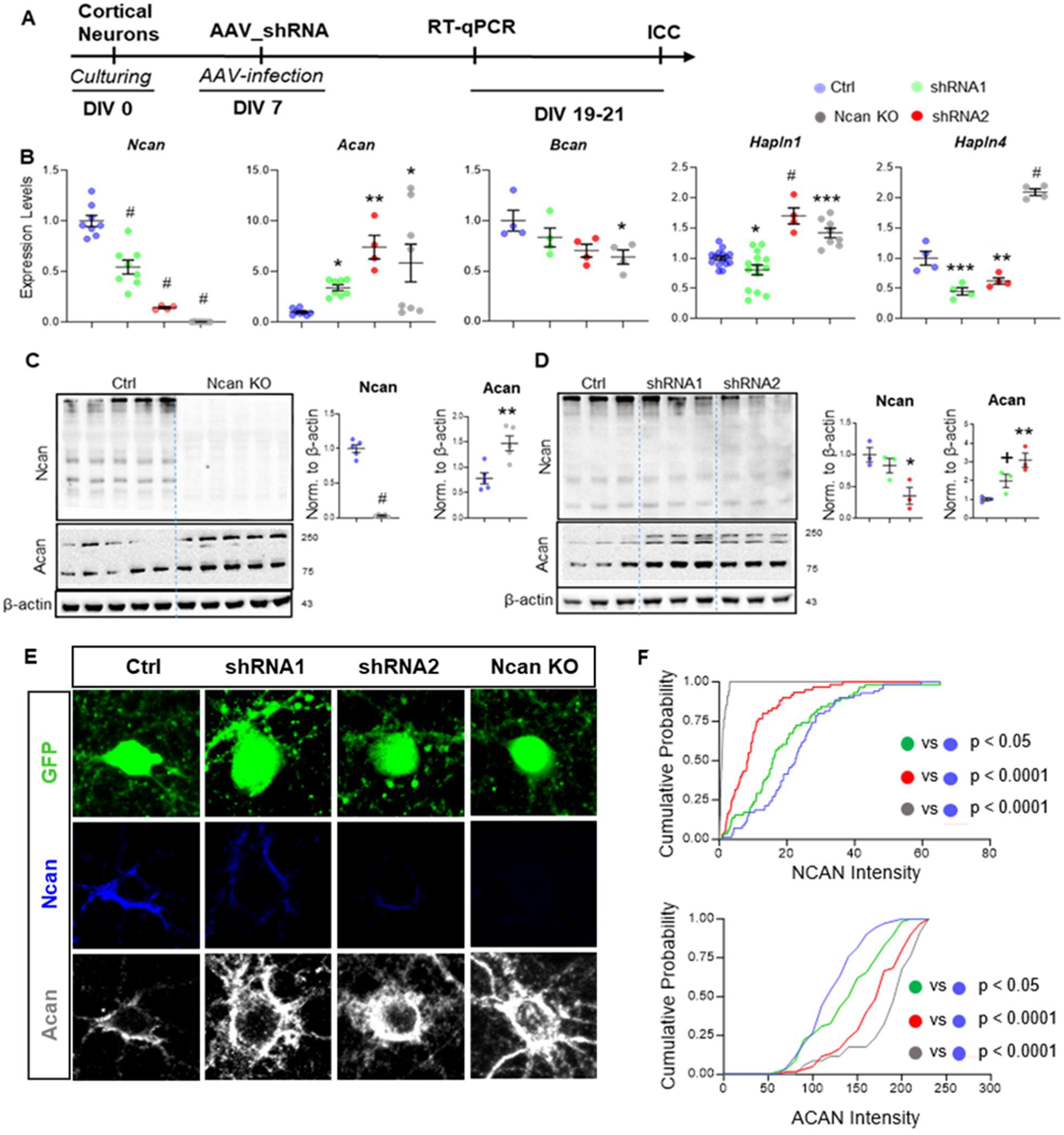
Effect of Ncan depletion on expression of key ECM components. **(A)** Timeline for Ncan knockdown experiments in primary cultured cortical cells. **(B)** qPCR analysis of ECM-associated gene products after Ncan depletion. **(C, D)** Western blot analysis of ECM proteins in Ncan KO and KD cortical cells, respectively. **(E, F)** Representative images of cortical cells infected with Ncan and control shRNA, expressing GFP, and associated analyses for Ncan and Acan intensities. The graphs show data from three independent experiments. One-way ANOVA with Sidak’s comparisons and Kolmogorov-Smirnov’s test were used for multiple comparison and the comparison of cumulative probability distributions, respectively. *p < 0.05, **p < 0.01, ***p < 0.001 and # p < 0.0001.

**Figure 2.**
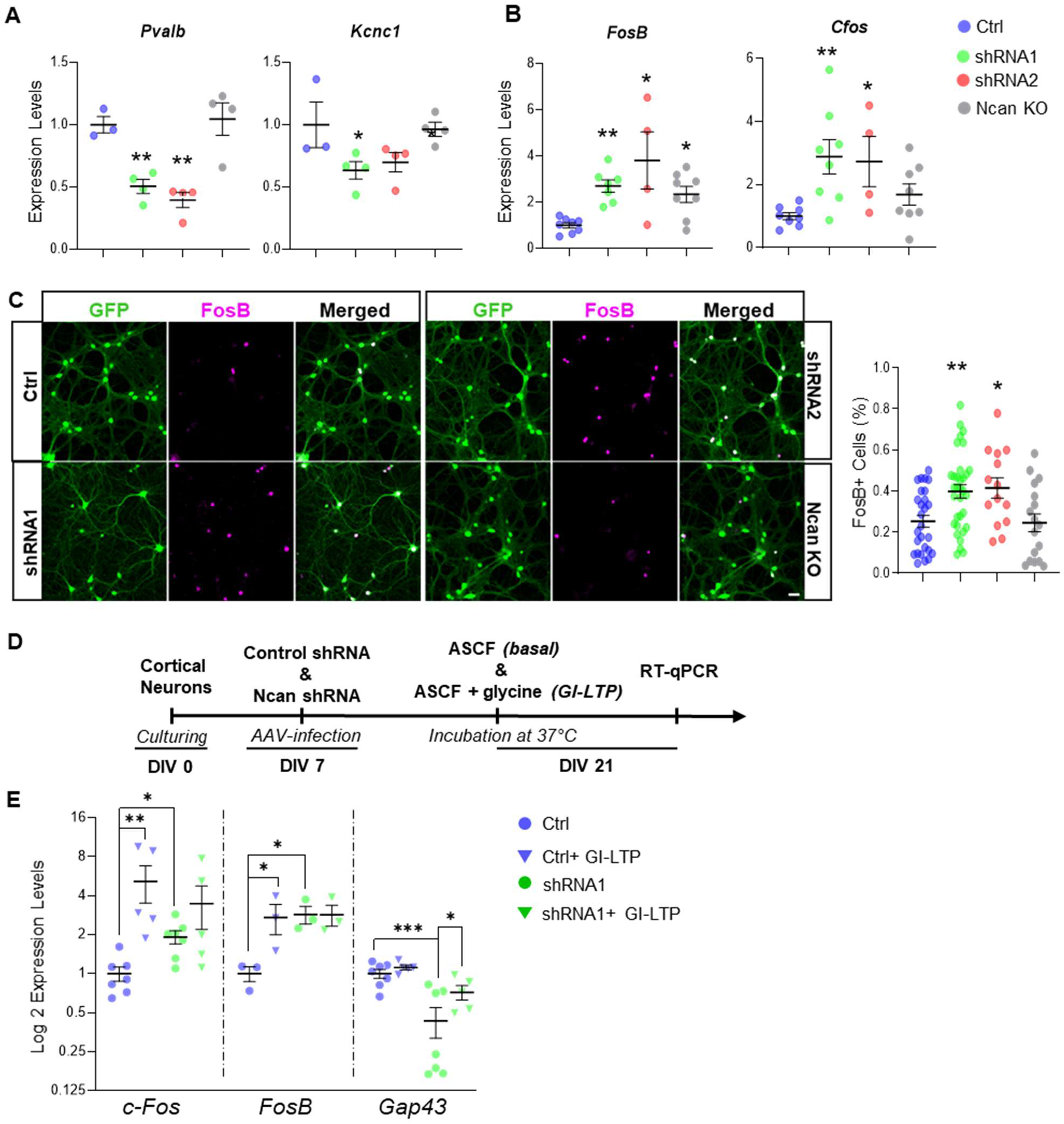
The depletion of Ncan results in compromised expression of inhibitory neuron markers and cellular activity indicators *in vitro*. **(A, B**) qPCR evaluation of inhibitory markers (*Pvalb* and *Kcnc1*) and cellular activity markers (FosB and cfos) after Ncan depletion. **(C)** Representative images of cortical cells infected with Ncan and control shRNA, expressing GFP, and associated analyses for the cell activity marker FosB. **(D)** Timeline for the GI-LTP experiment in cortical culture infected with control and shRNA1. **(E)** qPCR evaluation of activity-associated genes C-Fos, FosB and Gap43 in the basal and GI-LTP conditions. The bar graphs show the mean ± SEM from three independent experiments. one-way ANOVA with Sidak’s multiple comparisons test. *p < 0.05, **p < 0.01, and ***p < 0.001.

### Effects of Ncan knockdown on neuronal activity-dependent gene expression

Next, we investigated the effects of Ncan depletion on the expression of neuronal activity and synaptic plasticity-related genes. We used glycine-induced chemical long-term potentiation (GI-LTP), the global form of LTP, as it has been shown to mediate similar cellular processes as tetanus-induced LTP (Chen et al., 2011). Mature cortical cultures (DIV21) infected with Ncan shRNA and Ctrl at DIV7 were exposed to ACSF (basal condition) or ACSF supplemented with glycine to facilitate activation of NMDA receptors (GI-LTP condition) (Fig. 2D). As anticipated, GI-LTP stimulation resulted in an elevation of mRNA levels of *c-Fos* and *FosB* relative to baseline in Ctrl neurons. However, intriguingly, this effect was not observed in neurons after attenuation of Ncan expression (Fig. 2E), as there was a significant upregulation of the basal expression of *FosB* and *c- Fos* in shRNA1-expressing relative to Ctrl neurons, indicating an overall increased activity level (Fig. 2E). Then we investigated another synaptic plasticity-associated gene, the growth-associated protein 43 (GAP-43), which is involved in the presynaptic activity-dependent remodeling (Benowitz & Routtenberg, 1997) and axonal regeneration (Gil-Loyzaga et al., 2010). Interestingly, *Gap43* was strongly downregulated in shRNA-infected cortical neurons under basal conditions compared to Ctrl. GI-LTP enhanced the expression of *Gap43* in shRNA conditions, but not to the Ctrl levels (Fig. 2E). These findings suggest the important role of Ncan in regulation of neuronal activity and expression of GAP-43.

### Knockdown of Ncan expression enhanced frequency of synaptic events and bursts in vitro

Thus far, we provided evidence for an overall increase in neuronal activity after Ncan depletion in cortical neurons. To further confirm the role of Ncan in regulating the balance between excitatory and inhibitory inputs, we conducted whole-cell patch clamp recordings in cortical neurons treated with Ctrl and shRNA AAVs, and in Ncan KO neurons (Fig. 3A). Consistent with our expectations, we observed that Ncan depletion increased the synaptic inputs to cortical neurons infected with shRNA1 or shRNA2, and in Ncan KO neurons. In the Ncan shRNA1 or shRNA1 treated groups, all neurons displayed bursts of synaptic currents, whereas only 41.1% and 62.2% of the neurons in the Ctrl and Ncan KO exhibited such bursting phenotype (Fig. 3 B, C). Further analysis demonstrated similar increases in the burst frequency, duration, and number of events in the shRNA1 group and Ncan KO and less prominent changes in shRNA2 expressing neurons (Fig. 3D).

**Figure 3.**
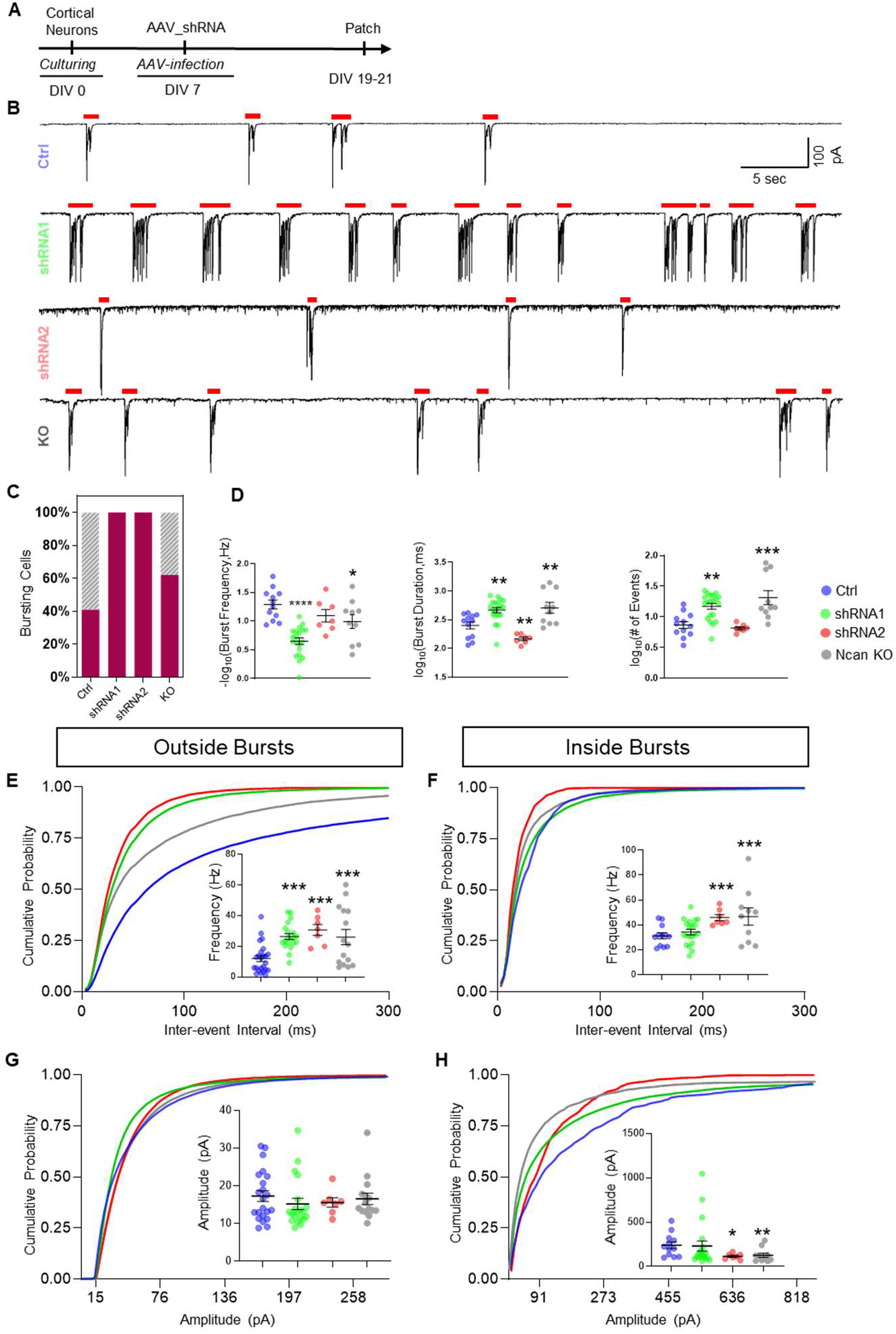
The depletion of Ncan leads to enhanced sEPSC and increased neuronal activity *in vitro*. **(A)** Timeline of patch-clamp experiments for spontaneous EPSC recording in cultured cortical cells. **(B)** Representative traces of spontaneous EPSCs recorded at −70 mV in the cortical neurons infected with control and shRNAs. Red lines above the traces indicate areas considered as a burst, according to designed method for burst detection. **(C)** Quantification of the proportion of cells exhibiting burst activity across experimental groups. **(D)** Quantification of three distinct electrophysiological parameters, burst frequency, burst duration, and number of spikes per burst, across multiple experimental groups. **(E, F)** Cumulative probability curves for inter-event intervals and calculated frequency in the regions of EPSC occurring within and outside of bursts. **(G, H)** Cumulative probability analysis of the amplitude in the regions of EPSC occurring within and outside of bursts. The bar graphs show the mean ± SEM from three independent experiments. One-way ANOVA with Sidak’s comparisons were used for multiple comparison. *p < 0.05, **p < 0.01, ***p < 0.001 and **** p < 0.0001.

Next, the recorded events were categorized into two distinct groups: burst-like events and events occurring between bursts (inter-burst events). Subsequently, we measured the frequency and amplitude of both types of events. Here, we compared burst features such as burst frequency, duration, and the number of events per burst, and found interesting patterns. Specifically, Ncan depletion was associated with a prominent increase in the frequency of spontaneous excitatory postsynaptic currents outside of bursts in shRNA-expressing and Ncan KO neurons as compared to Ctrl, indicating an elevated state of neuronal activity (Fig 3 E). The amplitude of sEPSCs was, on the other hand, comparable among all groups (Fig 3G). Intraburst analysis demonstrated a notable and significant increase in the frequency of events in the shRNA2 and Ncan KO groups (Fig. 3F). shRNA1-infected neurons exhibited a frequency of sEPSC similar to that observed in Ctrl neurons (Fig. 3G). Conversely, the amplitude of bursts was lower in shRNA2 and Ncan KO groups (Fig. 3H). Collectively, these findings strongly suggest that Ncan depletion contributes to elevation of neuronal activity.

### Knockdown of Ncan enhanced AnkG expression at AIS

Considering the accumulation of Ncan at the AIS, we further investigated the expression of AnkG, the major organizer of the AIS and regulator of neuronal excitability (Frischknecht et al., 2014) in Ncan-depleted or -deleted cultures. We plotted AnkG intensity relative to the distance from soma and fitted a simple exponential model to estimate various parameters of AnkG distribution including maximum AnkG intensity (Y_peak_), distance from the soma to the maximum intensity (X_peak_), the distance at which maximum intensity reached 50% (T_decay_), and AnkG intensity at the proximal part of the AIS (Y0) (Fig. 4A, B, C). For each experimental group, the goodness of fit (R²) exceeded the value of 0.9, indicating a high degree of model accuracy and that this model explains more than 90% of the inter-group variance. Our observations indicate that depletion of Ncan by shRNA1 and shRNA2 altered the localization of AnkG within the AIS of cortical neurons, resulting in a displacement of the maximal AnkG expression distance from the soma, as compared to the Ctrl group. In contrast, the localization of AnkG in the Ncan KO neurons remained unaffected (Fig. 4C, D). The maximum intensity of AnkG, i.e., Y_peak_, was found to increase in both shRNA1 and Ncan KO groups as compared to Ctrl, with no difference in shRNA2-expressing neurons (Fig. 4E). The distance at which AnkG expression reduces to 50%, i.e., Tau_decay_, showed significant differences between controls and all neurons with Ncan depletion or deletion, with shRNA1 and Ncan KO neurons showing similar values (Fig. 4F). No significant difference was observed in Y0 between the Ctrl and shRNA2 and shRNA1 while Ncan KO, on the other hand, showed a significant difference compared to the Ctrl (Fig. 4G). Analysis of the relationships between each of AnkG-dependent parameters and electrophysiological features across experimental groups revealed that expression of AnkG at soma and the peak values are proportional to log_10_ (# of events) (Fig. 4H, I), the location of the peak is inversely related to the burst duration (Fig. 4 J), while T_decay_ is related to the frequency of events outside of the bursts (Fig. 4K).

**Figure 4.**
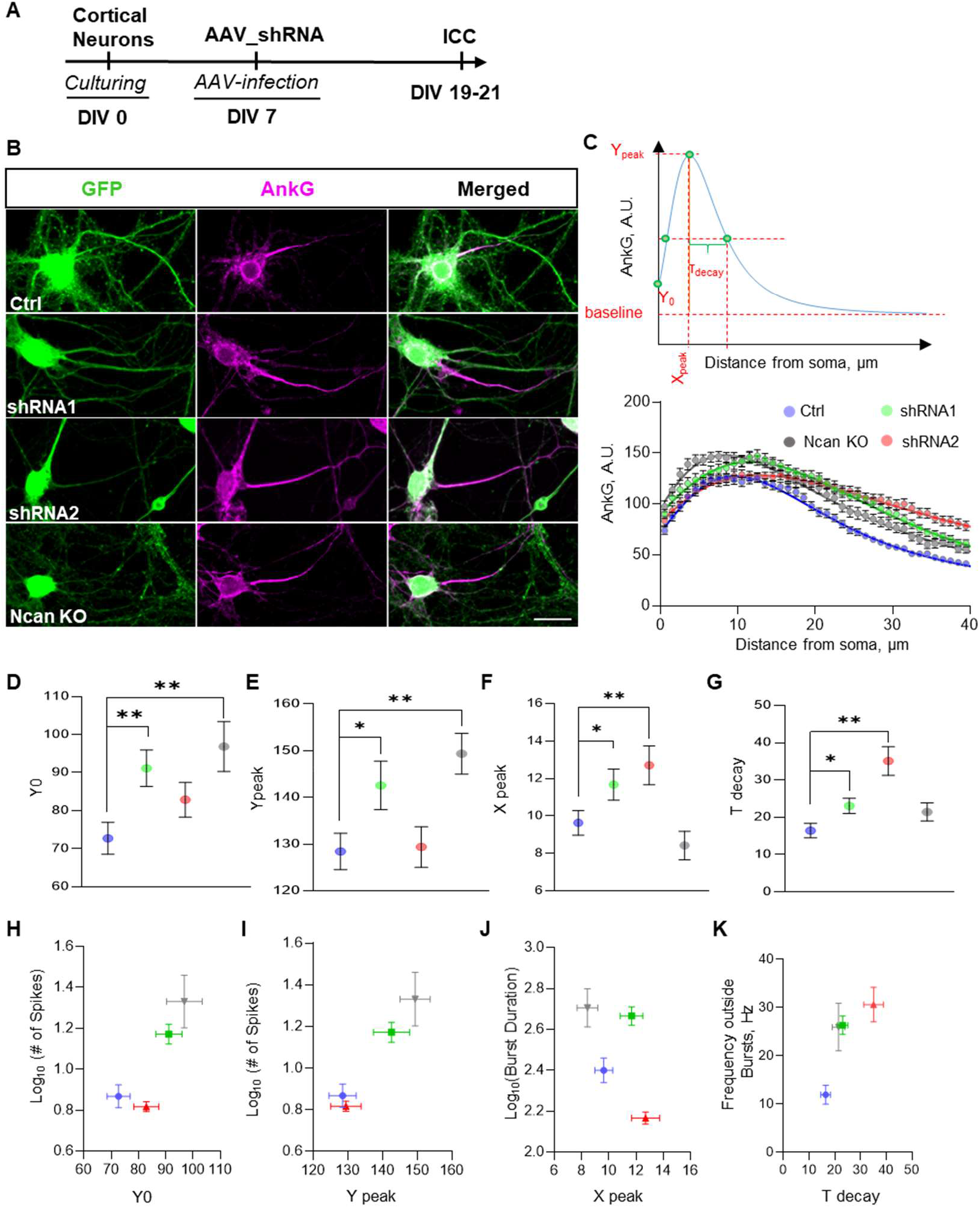
Ncan depletion alters AnkG distribution along the AIS. **(A)** Experimental timeline. **(B)** Representative images of cortical neurons infected with Ncan and control shRNA, expressing GFP, immunostained for AnkG. **(C)** Model depicting the quantification of various parameters related to AnkG expression, alongside the fitted model correlating with observed events. **(D, E, F, G)** Quantification of AnkG-associated parameters, including Y0 (the amount of Ankyrin at the proximal part of the axon), Ypeak (the peak height), Xpeak (distance from the soma to the maximum intensity), and Tdecay (the distance at which maximum intensity reached 50%), across experimental groups. **(H, I, J, K)** Correlation of various AnkG-dependent parameters with electrophysiological features across experimental groups. The bar graphs show the mean ± SEM three independent experiments. Statistical significance for panels D-G was calculated via bootstrap approach as described in Materials and Methods. *p < 0.05, and **p < 0.01.

In addition to this, immunoblot analysis showed no significant differences in the protein levels of Na1.2, Na1.6, and AnkG expression between Ctrl and Ncan KO neurons (Fig. 5A). While there was a tendency for downregulation in Nav1.2 in shRNA1 and a significant decrease in shRNA2 neurons, as well as for upregulation in Nav1.6 and AnkG expression in both shRNA groups (Fig. 5B). Western blot analysis of Ncan KO neurons infected with shRNA1 and shRNA2 revealed no significant difference in Acan, Bcan, Nav1.6, and AnkG protein levels, confirming the specificity of shRNA constructs (Fig. 5C).

**Figure 5.**
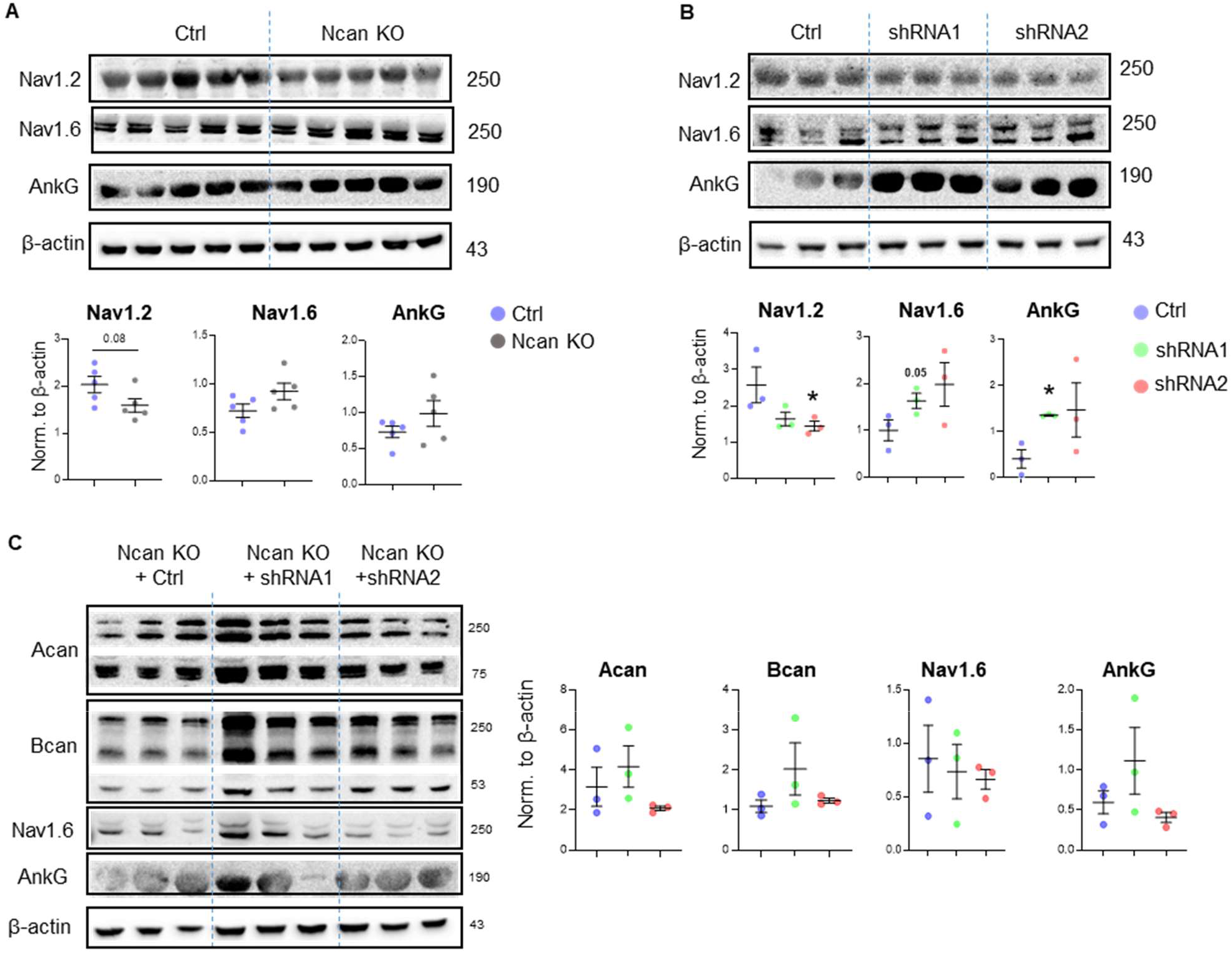
The depletion of Ncan changes the protein levels of sodium channels and AnkG. **(A)** Western blot analysis of Nav1.2, Nav1.6, AnkG, and β-actin from the cell lysates of wild-type and KO samples. **(B)** Western blot analysis of Nav1.2, Nav1.6, AnkG, and β-actin in cell lysates from control, shRNA1-and shRNA2-infected cultures. **(C)** Quantification of protein levels of Acan, Bcan, Nav1.6 and AnkG in KO cells infected with control, shRNA1 and shRNA2. The bar graphs show the mean ± SEM from three independent experiments. One-way ANOVA with Sidak’s multiple comparisons test. *p < 0.05.

### Knockdown of Ncan expression reorganized AnkG distribution at the AIS of both inhibitory and excitatory neurons

Finally, we assessed the impact of Ncan depletion on AnkG redistribution along the AIS of excitatory and inhibitory neurons separately. Here, we used GAD67 as a marker for identifying GABAergic neurons in the brain (Fig. 6A, B). After observing inconsistent effects on neuronal activity in cortical neurons infected with shRNA2, including extreme reduction of *Ncan* mRNA and upregulation of Bcan and Acan similar to shRNA1 and Ncan KO neurons, we chose to use shRNA1 for these experiments. It is possible that the contrasting effect on neuronal excitability in shRNA2 neurons was due to insufficient time for genetic compensation to occur (El-Brolosy & Stainier, 2017).

**Figure 6.**
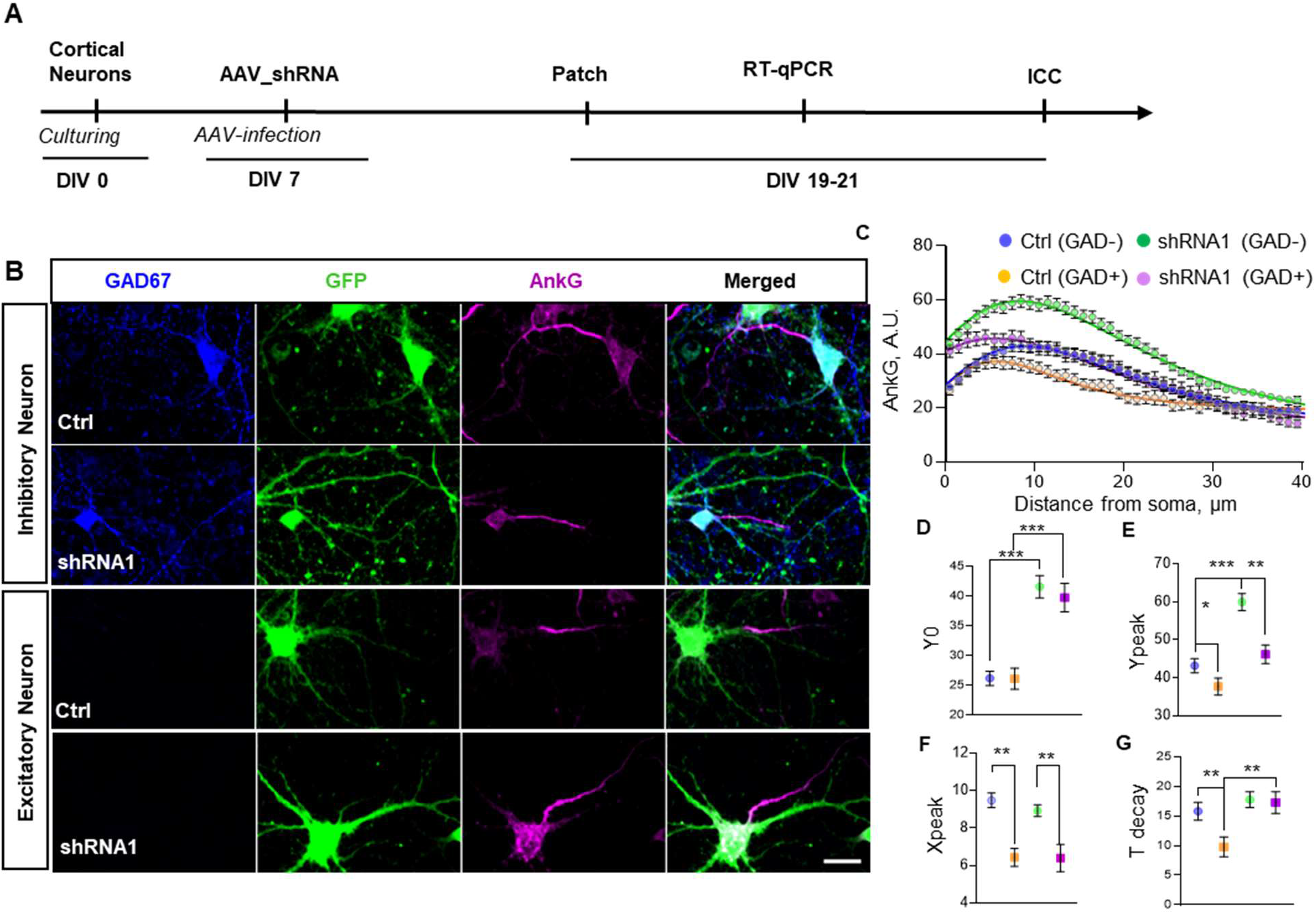
The depletion of Ncan alters AnkG parameters in excitatory and inhibitory neurons. **(A)** Experimental time-line. **(B)** Representative images of cortical neurons infected with Ncan and control shRNA, expressing GFP and immunostained for AnkG and the inhibitory neuron marker GAD67. **(C)** Graph depicting the fitted model correlating with observed AnkG expression in GAD67 positive and negative cells across experimental groups. **(D, E, F, G)** AnkG-dependent parameters in GAD67-positive and GAD67-negative neurons within control and shRNA1 experimental groups. The bar graphs show the mean ± SEM from three independent experiments. Statistical significance for panels D-G was calculated via bootstrap approach as described in Materials and Methods. **p < 0.01, and ***p < 0.001.

Our results show that the Y_0_ parameter, representing the AnkG intensity at the initial point of the AIS, is increased in Ncan shRNA1-expressing cells with no difference between GAD+ and GAD-neurons. The maximal intensity of AnkG was increased in GAD+ compared to GAD-neurons, and this effect appeared more prominent in Ncan shRNA1-expressing cells than in Ctrl (Fig. 6E). We noted an Ncan-independent proximal shift in the AIS of GAD+ inhibitory neurons in both Ctrl and Ncan shRNA1 conditions (Fig. 6F). Furthermore, the analysis revealed a reduction in T_decay_ in GAD+ relative to GAD-neurons within the Ctrl group (Fig. 6G). However, T_decay_ was increased by Ncan shRNA1 in GAD+ neurons, so no significant difference between GAD-and GAD+ cells was observed after the knockdown (Fig. 6G). Taken together, our findings reveal both common as well as neuron subtype-specific regulatory mechanisms of AIS organization, driven by Ncan.

## DISCUSSION

Our study demonstrates the importance of Ncan for expression of key organizing proteins in the PNNs and AIS in vitro, and highlights the Ncan role in regulation of neuronal activity with implications for synaptic plasticity.

### Ncan knockdown and the PNN molecular organization

Consistently, Ncan knockout and knockdown with two different shRNAs resulted in elevated expression of Acan at the mRNA and protein levels globally and specifically in the PNNs. Acan, being the primary and most abundant component of PNNs, plays a crucial role in providing structural stability to these nets. In the absence or reduction of Ncan, the upregulation of Acan may serve as a compensatory response to maintain the structural integrity and functional properties of PNNs (Ueno et al., 2018). Interestingly, a previous study has demonstrated a strong correlation between neuronal activity and *Acan* levels (Willis et al., 2022) after treating cortical neurons with bicuculline, an inhibitor of GABA-A receptors (Gribkoff et al., 2003; Johnston, 2013), thereby increasing the excitability of excitatory neurons. The increased neuronal activity upregulated c-Fos and Acan levels, and these genes returned to their control levels once silenced with tetrodotoxin (TTX) treatment (Willis et al., 2022). Similarly, we show that Ncan depletion upregulates neuronal activity, Acan and c-Fos levels. This, therefore, implies that the increase in Acan levels might be a consequence of elevated activity in Ncan-depleted cultures, which may represent a homeostatic protective response as it has been shown that PNNs support perisomatic inhibition and offer protective properties to fast-spiking interneurons (Cabungcal et al., 2013), and Acan is a central element in PNN formation (Giamanco et al., 2010; Rowlands et al., 2018).

For other ECM molecules, more complex dependences on Ncan were revealed. The level of Hapln1 mRNA was reduced by weak Ncan knockdown with shRNA1 and increased by strong knockdown with shRNA2 and by Ncan knockout. Hapln4 mRNA was decreased by both Ncan shRNAs, while Ncan knockout increased it. Thus, both Hapln1 and Hapln4 expression seem to follow a U-shaped expression pattern in relation to Ncan availability. These data show that Ncan regulates expression of aggrecan and link proteins as key organizers of PNNs, and that Ncan knockout results in some compensatory effects not observed after knockdown of Ncan.

### Ncan depletion increases neuronal activity

Ncan has been demonstrated to be crucial for the development of neuronal circuits and is known to be involved in controlling synaptic plasticity (Mohan et al., 2018; Zhou et al., 2001). Here, we found that Ncan knockdown resulted in an upregulation in activity-dependent gene expression of cFos and FosB, an increase in the number of cFos-immunopositive cells, and prominent enhancement in neuronal activity within and between bursts in cultured cortical neurons. One of underlying mechanisms seems to be mediated by the reduction in expression of PV and Kcnc1/K_v_3, the potassium channels important for fast-spiking of PV neurons. Previous studies have reported similar changes after PNN degradation by chondroitinase ABC (Willis et al., 2022; Lupori et al., 2023). The amount of parvalbumin in PV interneurons increases with their maturation (Bitzenhofer et al., 2020; Lupori et al., 2023) and has been shown to affect the acquisition of their mature phenotypes (Yang et al., 2014). Thus, it is plausible that Ncan knockdown shifts the excitation-inhibition ratio towards excitation, modulating activity of PV cells. A critical role for the ECM in modulating neuronal firing activity and maintaining the delicate balance between excitation and inhibition has been well documented in previous studies (Gottschling et al., 2019; Hayani et al., 2018; Lensjo et al., 2017). Previous findings suggested that the ECM aids in preserving the balance between excitatory and inhibitory neurons by upholding inhibitory connections (Saghatelyan et al., 2001, 2004). Consequently, the removal of ECM resulted in a significant decrease in inhibitory synapse density (Dzyubenko et al., 2021). Consistent with our findings, a recent investigation proposed that astrocyte-secreted Ncan has a critical function in preserving the integrity of inhibitory synapses. The Eroglu lab showed in a preprint that Ncan deletion reduces the frequency of miniature inhibitory postsynaptic currents (mIPSCs) in cortical neurons of Ncan KO mice (Irala et al., 2023).

### Ncan knockdown and AIS remodeling

The precise positioning and composition of the AIS at the proximal axon play a critical role in determining the excitability of neurons (Huang & Rasband, 2018). Here, we observed a distal shift in the peak of AnkG expression following Ncan depletion in shRNA-infected neurons. No such changes were found in Ncan KO neurons. The shift correlated with increased neuronal excitability. This finding aligns with a theoretical analysis that emphasizes the increase in neuronal excitability due to a distal shift in the AIS in the absence of hyperpolarizing currents (Goethals & Brette, 2020). AIS exhibits a high concentration of voltage-gated ion channels, particularly sodium channels. Interestingly, in line with our observation of reduced Na_v_1.2 concentration after Ncan depletion, Zhang and colleagues showed that deficiency in Nav1.2 sodium channels induces neuronal hyperexcitability in adult mice (Zhang et al., 2021). The result provides a potential explanation for the clinical observation that individuals with Na_v_1.2 deficiency experience seizure activity (Nadella et al., 2022). Alterations in the expression and functionality of Na_v_1.6 sodium channels have a significant impact on the firing properties of neurons (Zybura et al., 2021), and effective elimination of action potential propagation has been observed in Na_v_1.6 KO mice (Ye et al., 2018), as well as after pharmacological inhibition (Johnson et al., 2022). A comparable pattern of neuronal activity was observed in our experiments when Ncan depletion resulted in the upregulation of Na_v_1.6 at the protein level, leading to neuronal hyperexcitability. Although the exact role of Ncan in redistribution of AnkG is unclear at this stage, it is in line with previous studies that suggested a role of the ECM in the organization and functionality of the AIS via interactions with NF186 and sodium channels (Srinivasan et al., 1998; Hedstrom et al., 2007; Susuki et al., 2013).

### Ncan knockdown and synaptic plasticity

To further study the effects of Ncan downregulation on synaptic plasticity, we searched for expression changes of activity-related genes after induction of chemical LTP. One major finding is the reduced expression of *Gap-43* in both basal and GI-LTP conditions. Previous studies have shown a strong relationship between *Gap-43* expression and memory. For example, overexpression of *Gap-43*, as seen in the hippocampus of Alzheimer’s patients, impaired learning and memory whereas moderate expression improved it (Holahan et al., 2007; Rekart et al., 2005). Also a reduced expression of *Gap-43* as found here resulted in memory dysfunction (Rekart et al., 2005). On the other hand, the enhanced basal expression of Fos-B observed after Ncan knockdown in cortical neurons is also an interesting finding as its expression strongly correlates with neural epileptiform activity (Vossel et al., 2016). In AD patients and several AD animal models, subclinical epileptiform activity has been reported, which correlates strongly with the chronic accumulation of *Fos-B*, impacting also the expression of *c-Fos* (Corbett et al., 2017; Kam et al., 2016). Interestingly, overexpression of *Fos-B* is also shown to impact hippocampal learning and memory, possibly through *Fos-B*-dependent induction of immature spines (Eagle et al., 2015). Moreover, the acute neuronal activity-dependent upregulation of *c-Fos* that is essential for LTP-dependent synaptic plasticity (Calais et al., 2013), i.e. the cellular and molecular form of learning and memory (Flavell & Greenberg, 2008), did not occur in cortical cells after Ncan knockdown. This is in agreement with a previous work in which basal *c-Fos* expression was found to be upregulated in memory-impaired aged mice (Haberman et al., 2017).

### Potential relation to human neuropsychiatric disorders

Genome-wide association studies (GWAS) have previously identified a polymorphism in the human NCAN gene, rs1064395, in the risk for bipolar disorder (Cichon et al., 2011), and for schizophrenia (Mühleisen et al., 2012). *Post mortem* eQTL analyses have suggested that the NCAN polymorphism might be associated with slightly increased levels of Ncan and the neighboring gene product HAPLN4 in the dorsolateral PFC (Assmann et al., 2021), while the association of the risk variant with the manic phenotype resembles the behavioral phenotype of NCAN-deficient mice (Miro et al., 2012), pointing to a potentially detrimental effect of both increased as well as decreased Ncan levels.

Our present findings have revealed a regulatory role for Ncan in the expression and cellular distribution of ankyrin G. The ANK3 gene, which encodes the human ankyrin G protein, has also been identified in GWAS of schizophrenia (Schizophrenia Psychiatric GWAS Consortium, 2011) and bipolar disorder (Ferreira et al., 2008; Schulze et al., 2009). Notably, expression studies suggest an increased expression of ANK3 in both disorders (Wirgenes et al., 2014), mirroring the increased ankyrin G expression upon Ncan depletion observed here. Furthermore, *post mortem* investigations have revealed increased expression of K_v_3.1-containing potassium channels in parvalbumin-positive GABAergic neurons in patients with schizophrenia, which normalize after treatment with neuroleptics (Yanagi et al., 2014). That finding is to some extent mirrored by the increased expression of Kcnc1 in Ncan-deficient parvalbumin-positive neurons.

More broadly, our results suggest that reduced Ncan expression might contribute to an altered excitation-inhibition ratio hypothesized to be a key circuit mechanism in the major psychoses (Howes & Shatalina, 2022; Grent-’t-Jong et al., 2023), which, at a systems level, may be reflected by a manic phenotype (Miro et al., 2012) and tonically increased hippocampal activity (Bartsch et al., 2023; Assmann et al., 2021).

## Conclusion

In conclusion, our study highlights the importance of Ncan in regulating the expression patterns of PV, Acan, Bcan and AnkG, neuronal activity and synaptic plasticity in cortical neurons. Our results point to a complex regulatory role of Ncan in the regulation of multiple proteins implicated in neuropsychiatric disorders like bipolar disorder and schizophrenia, thereby shedding new light on the previously reported genetic association of the human NCAN gene with these disorders.

## Disclosure statement

Authors declare no conflicts of interests.

## Acknowledgments

We thank Katrin Boehm and Jenny Schneeberg for technical support. This work has been supported by DAAD (fellowship to D.B.A), Deutsche Forschungsgemeinschaft (DFG, German Research Foundation) – Project-ID 425899996 – SFB 1436 (to A.D., B.H.S. and C.S). and RTG 2413 SynAge (to A.D. and C.S).

## Author’s contribution

D.B.A produced all AAVs, plated and infected cortical neurons. D.B.A also, performed all ICC and RT-qPCR; H.M performed all electrophysiological recordings and WB analysis. S.A. performed analysis of electrophysiological data and parameters of AnkG distributions. C.S. supervised breeding of Ncan KO mice and biochemical analyses; R.K and S.A designed Ncan shRNA AAVs. D.B.A wrote all Fiji scripts for image analysis of all samples. D.B.A and H.M. wrote the manuscript. A.D. with D.B.A designed the study and A.D. supervised all data analysis. All authors contributed to the editing of the manuscript.

---

**Supplementary Table 1.**
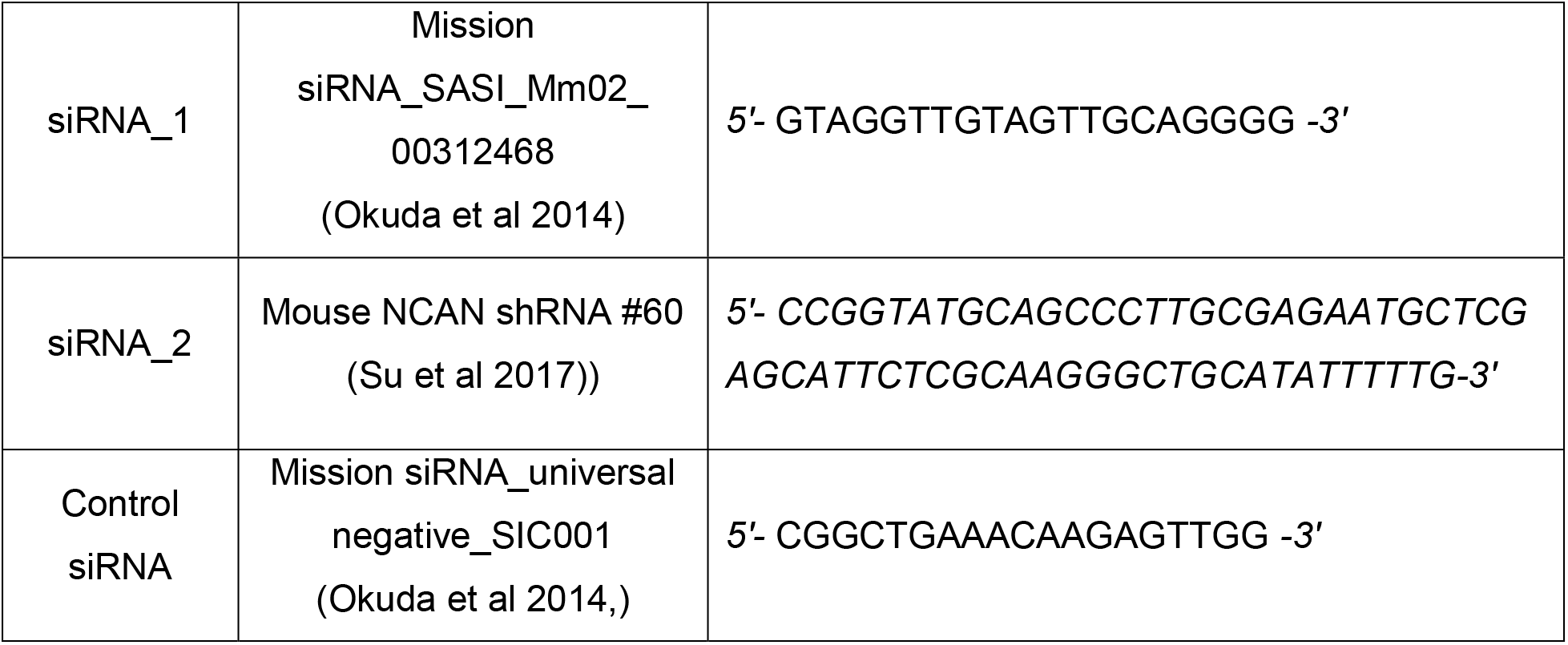
siRNA sequences used in this study.

**Supplementary Table 2.**
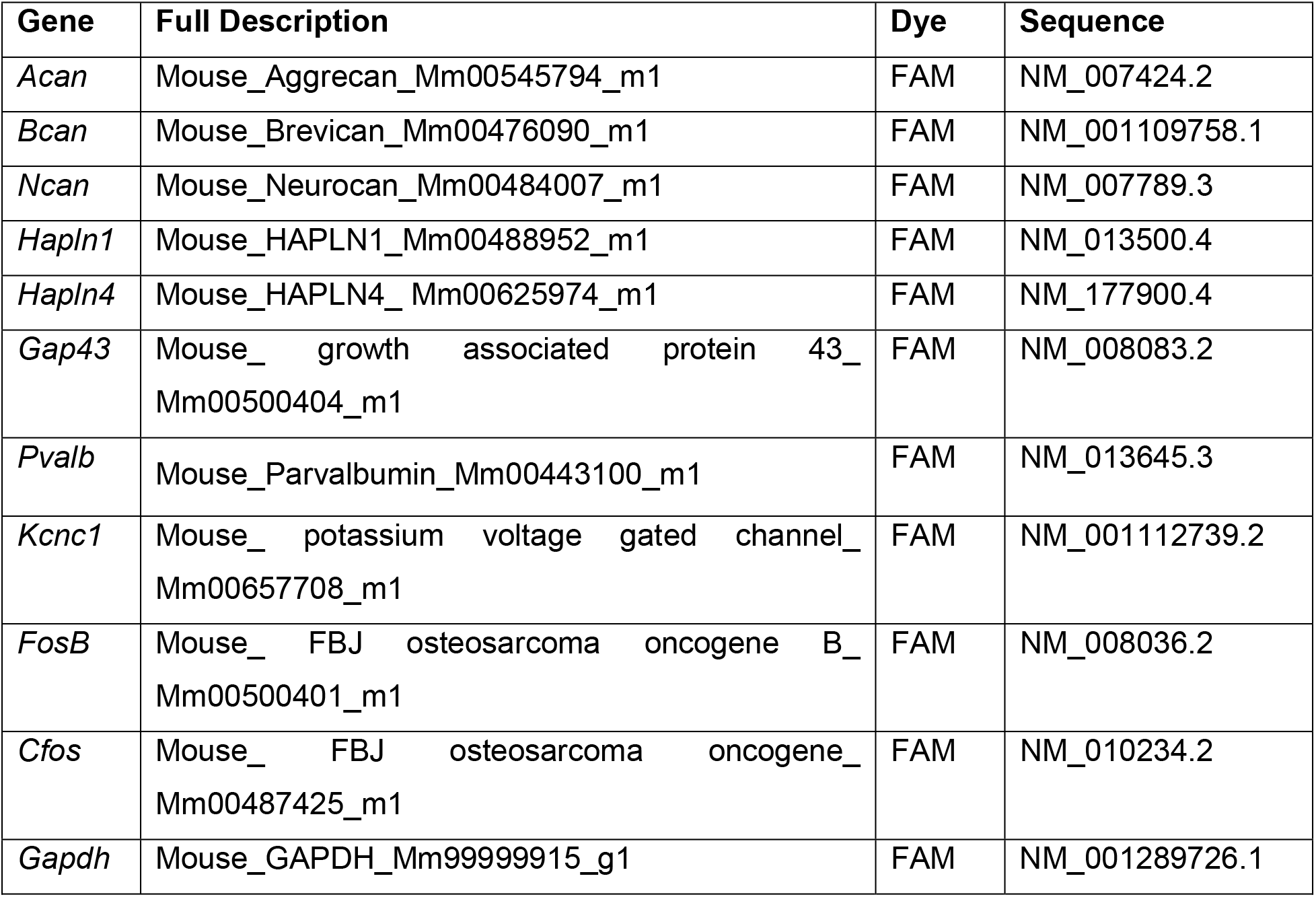
Taqman probes used for real-time PCR analysis.

**Supplementary Table 3.** Primary antibodies and their dilutions for ICC and WB.

| Antibody | Dilutions | Species | Provider/Cat. No. |
| --- | --- | --- | --- |
| Acan | 1:500 | Rabbit | Millipore; AB1031 |
| Bcan | 1:1000 | Guinea pig | LIN Magdeburg (John et al., 2006) |
| Ncan | 1:100 | Sheep | Synaptic systems; AF5800 |
| FosB | 1:500 | Rabbit | Cell signaling; 2251 |
| AnkG | 1:500 | Guinea pig | Synaptic systems; 386004 |
| Nav 1.2 | 1:500 | Rabbit | Alomone Labs; ASC002 |
| Nav 1.6 | 1:500 | Rabbit | Alomone Labs; ASC009 |
| Gad67 | 1:1000 | Mouse | Merck; MAB5406 |

**Supplementary Table 4.** Secondary antibodies and their dilutions for ICC.

| Antibody<br>(Alexa Flour) | Dilution | Species | Provider |
| --- | --- | --- | --- |
| Anti-Mouse | 1:1000 | Goat | Invitrogen |
| Anti-Guinea pig | 1:1000 | Goat | Invitrogen |
| Anti-Rabbit | 1:1000 | Goat | Invitrogen |
| Anti-Sheep | 1:1000 | Donkey | Invitrogen |

## Notes

### Competing Interest Statement

The authors have declared no competing interest.

